# Ultra-high field (7T) functional magnetic resonance imaging in amyotrophic lateral sclerosis: a pilot study

**DOI:** 10.1101/2020.12.02.408278

**Authors:** Robert L. Barry, Suma Babu, Sheeba Arnold Anteraper, Christina Triantafyllou, Boris Keil, Olivia E. Rowe, D. Rangaprakash, Sabrina Paganoni, Robert Lawson, Christina Dheel, Paul M. Cernasov, Bruce R. Rosen, Eva-Maria Ratai, Nazem Atassi

## Abstract

Amyotrophic lateral sclerosis (ALS) is a neurodegenerative disease of the central nervous system that results in a progressive loss of motor function and ultimately death. It is critical, yet also challenging, to develop non-invasive biomarkers to identify, localize, measure and/or track biological mechanisms implicated in ALS. Such biomarkers may also provide clues to identify potential molecular targets for future therapeutic trials. Herein we report on a pilot study involving twelve participants with ALS and nine age-matched healthy controls who underwent high-resolution resting state functional magnetic resonance imaging at an ultra-high field of 7 Tesla. A group-level whole-brain analysis revealed a disruption in long-range functional connectivity between the superior sensorimotor cortex (in the precentral gyrus) and bilateral cerebellar lobule VI. *Post hoc* analyses using atlas-derived left and right cerebellar lobule VI revealed decreased functional connectivity in ALS participants that predominantly mapped to bilateral postcentral and precentral gyri. Cerebellar lobule VI is a transition zone between anterior motor networks and posterior non-motor networks in the cerebellum, and is associated with a wide range of key functions including complex motor and cognitive processing tasks. Our observation of the involvement of cerebellar lobule VI adds to the growing number of studies implicating the cerebellum in ALS. Future avenues of scientific investigation should consider how high-resolution imaging at 7T may be leveraged to visualize differences in functional connectivity disturbances in various genotypes and phenotypes of ALS along the ALS-frontotemporal dementia spectrum.

## 1 Introduction

Amyotrophic lateral sclerosis (ALS) is characterized by the progressive degeneration of motor neurons and their axonal connections in the brain and spinal cord. The disease presents as a continuous loss of motor functions and ultimately results in death. ALS is known to be associated in many individuals with a spectrum of extra-motor dysfunction including behavioral changes, cognitive impairment, dementia and mood changes ^1^. Although rare, there are growing observations that some individuals with “ALS plus” syndromes may have other associated neurological features including extrapyramidal/parkinsonian features, diminished taste/smell, cerebellar ataxia, parasomnias, or eye gaze abnormalities ^1^– ^8^. This suggests that ALS may also be viewed as an overlapping neurodegenerative disorder with widespread involvement of different brain regions.

From a neuropathological standpoint, ALS is recognized as a TDP43 proteinopathy in 98% of all cases (except forms of ALS with SOD1 and FUS gene expressions) ^9^. The Braak staging of spreading TDP43 inclusion pathology burden in a large postmortem ALS cohort suggests that the disease pathology extends beyond the well known motor cortical system in later stages of disease to also include precerebellar nuclei, prefrontal cortex, and the striatum^10^. There is an unmet need to expand ongoing *in vivo* clinical research to evaluate reliable and sensitive neuroimaging biomarkers ^11^ that can identify and map neural networks that degenerate with ALS and more importantly their relevance and predictability to clinical disability progression ^12^–^20^. There has also been growth in multimodal neuroimaging techniques in the past decade, which provide unique opportunities to leap forward toward developing novel and quantitative biomarkers of disease localization, tracking disease progression, and improving phenotypic classifications in ALS ^21^–^23^.

At 1.5 or 3 Tesla, diffusion magnetic resonance imaging (MRI) studies of ALS have revealed widespread structural damage across motor and non-motor regions^24^–^38^. The role of functional MRI (fMRI) in the study of neurodegenerative disorders continues to expand^39^ and maps brain networks based upon the temporal coherency of hemodynamic responses in a ‘resting state’ or evoked via one or more tasks. Prior fMRI studies at 1.5 or 3T (and one at 4.7T) have revealed the impact of disease on brain functions ^40–45,26,28,46–53^ and modulation of functional ^54,55,28,33,34,56,57,36–38^ or effective ^58^ connectivity. The literature on resting state fMRI (rs-fMRI) in ALS is relatively sparse, and there is minimal convergence of findings across different cross-sectional and longitudinal cohorts ^59^. However, several rs-fMRI studies have noted aberrations (increases or decreases) in functional connectivity in ALS participants involving the sensorimotor network (SMN) ^54^,^28^,^56^,^33^,^36^,^38^ as well as other intrinsic networks ^60^.

To date, only a handful of 7 Tesla (7T) ALS studies have been published in the brain ^61^–^67^ and spinal cord ^68^. The benefits of imaging at an ultra-high field of 7T include increases in spatial signal-to-noise and contrast-to-noise ratios, functional contrast-to-noise ratio, and spectral dispersion ^69^–^72^. In the cervical cord, high-resolution T_2_*-weighted images demonstrated sharp delineation between gray and white matter, and white matter T_2_* hyperintensities were visualized in regions of ALS pathology ^68^. In the brain, a previous 7T ALS study reported hyperintensities in R_2_* maps in the motor cortex of ALS participants, which correlated with iron accumulation in *post mortem* pathological studies^62^. Anatomical imaging of the deep layers of the primary motor cortex at 7T revealed atrophy and T_2_* hypointensities in participants with ALS that correlated with upper motor neuron (UMN) impairment and disease progression rate^63^, and increases in magnetic susceptibility measured via quantitative susceptibility mapping co-localized with T_2_* hypointensities in the middle and deep layers ^64^. Diffusion imaging at 7T did not reveal whole-brain differences between controls and ALS participants, but analyses of the corticospinal tracts (CSTs) revealed decreased fractional anisotrophy and increased magnetization transfer ratio in ALS participants^65^. Single voxel magnetic resonance spectroscopy (MRS) of the left precentral gyrus at 7T revealed decreases in *N* -acetylaspartate (NAA), glutamate, and total NAA (*N* -acetylaspartate+*N* -acetylaspartylglutamate) in ALS participants, which correlated with forced vital capacity; additionally, the ratio of total NAA to total creatine (creatine+phosphocreatine) correlated with overall functional decline as measured by decreasing ALS Functional Rating Scale-Revised (ALSFRS-R) ^66^ scores. Another 7T MRS study reported a lower ratio of total NAA to *myo*-inositol in both the motor cortex and pons of ALS participants, and that levels of total NAA, *myo*-inositol, and glutamate in the motor cortex of ALS participants were influenced by the extent of disease progression^67^.

We are not aware of previous reports studying ALS using 7T fMRI. Therefore, the pilot study presented herein contributes to the existing body of 7T work by investigating group-level differences in functional networks between healthy controls and ALS participants using high-resolution rs-fMRI with spatial coverage extending from the cerebrum to the cerebellum. We additionally assessed the association between significant imaging findings and ALS clinical variables.

## 2 Material and methods

### 2.1 Study cohort

This cross-sectional pilot study was conducted at the Athinoula A. Martinos Center for Biomedical Imaging, Massachusetts General Hospital, Charlestown, Massachusetts, USA. Scans were performed within a 4-year period between 2011 and 2015. Subjects included in the current study (*N* = 21) were drawn from a cohort of 25 subjects previously reported by our group ^66^ who provided a high-resolution rs-fMRI scan upon completion of the main MRS protocol. All subjects provided written, informed consent through a protocol approved by the Partners Human Research Committee.

Twelve participants who met the revised El Escorial criteria for at least possible ALS and nine age-matched healthy controls were enrolled in this study. All ALS participants met the institutional MRI safety criteria and were able to tolerate lying flat for the scan duration. All participants had to have no other neurodegenerative disease diagnosis. Baseline characteristics of the study cohorts are detailed in Table 1. Family history of ALS was not collected for these participants. One ALS participant was confirmed to have C9orf72 repeat expansion. ALS causative genetic mutations remained unknown for the remaining 11 participants. All ALS participants underwent standard clinical outcome assessments collected by certified raters at a matching time point to the scan including ALSFRS-R^73^, slow vital capacity (SVC), and a quantitative muscle strength test using hand-held dynamometry.

**Table 1:** Baseline characteristics for healthy controls and participants with ALS.

| Baseline characteristic | ALS ($n = 12$)<br>%($n$) / $\mu(\sigma)$ | Controls ($n = 9$)<br>%($n$) / $\mu(\sigma)$ |
| --- | --- | --- |
| Age at screening (yrs) | 56.8 (10.3) | 53.2 (11.1) |
| Genetic abnormality | unknown (11)<br>C9orf72 positive (1) | n/a |
| Male | 75% (9) | 55% (5) |
| Caucasian | 100% (12) | 100% (9) |
| Symptom onset to diagnosis (months) | 10.3 (7.0) | n/a |
| Symptom onset to scan (months) | 30.3 (23.4) | n/a |
| Limb onset | 67% (8) | n/a |
| Baseline SVC (% predicted) | 93.4% (22.2%) | n/a |
| ALSFRS-R at baseline | 38.1 (4.8) | n/a |
| Estimated ALSFRS-R slope<br>pre-baseline ((48 – baseline<br>ALSFRS-R) / disease duration<br>(points/month) | 0.46 (0.35) | n/a |
| Revised El Escorial criteria | definite (5)<br>probable (3)<br>probable lab supported (1)<br>possible (3) | n/a |
| Riluzole use | 75% (9) | n/a |

### 2.2 Data acquisition

Experiments were performed on a whole-body 7 Tesla system (Siemens Healthineers, Erlangen, Germany) using a single-channel transmit birdcage volume coil and a custom-built 32-channel receive coil ^74^.

All subjects completed a 45-min brain imaging session without contrast, which included a high-resolution T1-weighted anatomical scan, MRS (previously reported ^66^), and a high-resolution rs-fMRI scan. High-resolution anatomical images were acquired with the following parameters: field of view (FOV) = 256 mm× 256 mm× 176 mm, voxel size = 1 ×1× 1 mm^3^, TE = 1.48 ms, inversion time = 1100 ms, repetition time (TR) = 2530 ms, flip angle = 7°. High-resolution blood oxygenation level dependent (BOLD) functional images were acquired with the following parameters: FOV = 190 mm×190 mm×108 mm, voxel size = 1.15 *×* 1.15 *×* 1.18 mm^3^, TE = 22.6 ms, TR = 5000 ms (except one subject where TR = 5560 ms inadvertently), flip angle = 84°, number of volumes = 94 (scan time = 7.8 min). Subjects were instructed to remain still and in a state of wakeful rest prior to the functional scan. Reported fMRI parameters are the mean across subjects due to minor adjustments to the protocol over the course of the 4-year study, and subject-specific adjustments to obviate specific absorption rate limits and peripheral nerve stimulation.

### 2.3 Data processing and analysis

Data were processed and analyzed using the CONN toolbox ^75^ release 18.b. Structural scans were translated to (0,0,0) coordinates and normalized to Montreal Neurological Institute (MNI) space ^76^ with simultaneous segmentations of gray matter, white matter, and cerebrospinal fluid (CSF). Functional scans were preprocessed using CONN’s recommended pipeline for volume-based analyses: i) realignment and unwarping (including rigid-body motion estimation and correction), ii) centering to (0,0,0) coordinates, iii) slice-timing correction, iv) ARtifact detection Tools (ART)-based outlier detection for data ‘scrubbing’ (www.nitrc.org/projects/artifact_detect) with thresholds of ±3*σ* for deviation from global mean and 1.5 mm for framewise displacement, v) normalization to MNI space with segmentations of tissues and CSF, and vi) spatial smoothing using an 8-mm full-width-at-half-maximum Gaussian kernel ^77^ to reduce between-subject anatomical variability in preparation for group analyses. Physiological noise regressors included five principal components of anatomically-derived white matter and CSF segments’ time series ^78^, respectively, the six estimated rigid-body motion parameters and their first derivatives, and ART-derived outliers. Additional preprocessing steps included linear detrending and band-pass filtering (0.008–0.09 Hz).

An exploratory second-level analysis of functional connectivity was performed using CONN’s seed-to-voxel analysis tool that considers numerous seed regions throughout the brain. Each seed was a 2 × 2 × 2 mm^3^ region centered at a given MNI coordinate. A functional contrast of Controls *>* ALS participants was selected to probe group-level differences between controls and participants with ALS. Results from this contrast (presented hereafter) were saved and projected onto a spatially unbiased atlas of the cerebellum using the spatially unbiased infra-tentorial template (SUIT) version 3.0^79–82^. *Post hoc* seed-to-voxel analyses with the same contrast were then performed using three atlas (AAL)-derived regions: left cerebellar lobule VI, right cerebellar lobule VI, and combined (left and right) cerebellar lobule VI.

## 3 Results

The cohorts of ALS participants (*n* = 12) and controls (*n* = 9) did not differ significantly in age or sex (*p* = 0.39 and *p* = 0.38, respectively, using Wilcoxon rank sum tests). The aggregate rigid-body motion parameters were also not significantly different between groups (*p* = 0.66, *t* -test), which is an important quality assurance observation to ensure that group-level differences in functional connectivity were not erroneously driven by group-dependent motion ^83^. The number of data points ‘scrubbed’ between groups was also not significantly different (median ± median absolute deviation: 5 ± 3.85 for controls vs. 5 ± 6.61 for ALS participants; *p* = 0.83 using a Wilcoxon rank sum test). No subjects were excluded due to excessive motion.

Across all regions considered in this exploratory analysis (voxel threshold: *p <* 0.005 uncorrected, two-sided; cluster threshold: *p <* 0.05 with false discovery rate correction), we observed a decrease in functional connectivity between the superior sensorimotor cortex (MNI coordinates = (0, –31, 67), which anatomically localizes to the inter-hemispheric fissure and in close proximity to the proximal leg representation area of the motor cortex) and lobule VI of the cerebellum in ALS participants vs. matched controls. Figure 1a presents the functional contrast Controls *>* ALS participants overlaid onto a single 2D axial slice through the mid-cerebellum. While a bilateral region of the cerebellum exhibited higher connectivity in controls, no regions exhibited higher connectivity in ALS participants at the same threshold. (Additional *post hoc* analyses within groups showed that these results were due to an absence of significant correlations in ALS participants.) Figure 1b projects these results onto a cerebellar flatmap ^82^. The group-level results are remarkably symmetric and predominantly mapped onto bilateral cerebellar lobule VI (involving both medial and lateral aspects of this cerebellar lobule).

**Figure 1:**
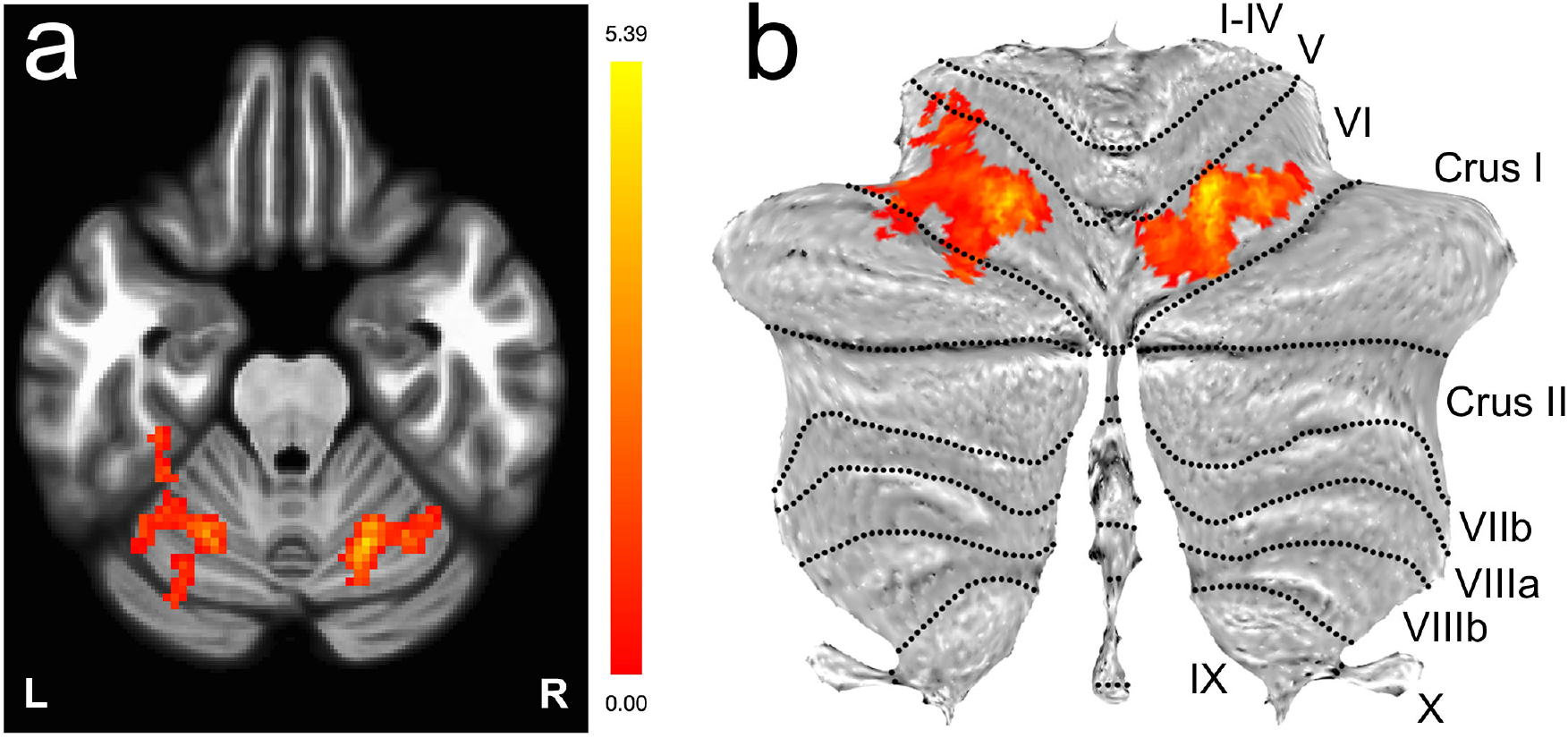
Decreased functional connectivity between the sensorimotor cortex and bilateral lobule VI of the cerebellum visualized via the functional contrast Controls *>* ALS participants (*p <* 0.005 uncorrected, two-sided; cluster threshold: *p <* 0.05 with false discovery rate correction) for a seed region located in the superior sensorimotor cortex (MNI coordinates = (0, –31, 67)). The results are presented as a) an overlay onto a single 2D axial slice through the cerebellum, and b) a projection onto a cerebellar flatmap^82^

Figure 2 visualizes results from three *post hoc* analyses using cerebellar lobule VI seed regions, presented as superior views of the partially-inflated cortical surface (voxel threshold: *p <* 0.001 uncorrected, two-sided; cluster threshold: *p <* 0.05 with false discovery rate correction). Group-level differences between ALS participants and controls predominantly mapped to the postcentral and precentral gyri with secondary differences in the superior parietal lobules and the precuneus cortex. Detailed results are presented in Table S1.

**Figure 2:**
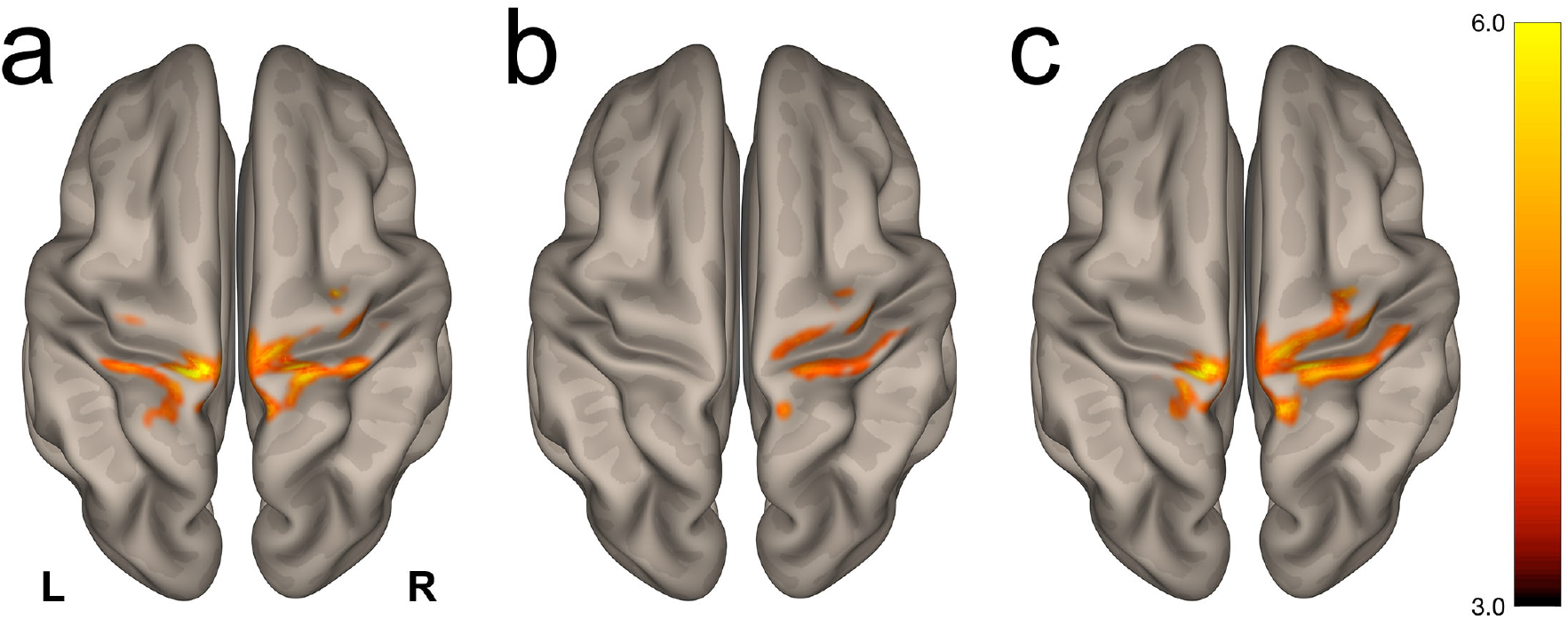
Visualization of the *post hoc* seed-to-voxel contrast Controls *>* ALS participants using atlas-derived a) left, b) right, and c) bilateral cerebellar lobule VI as seed regions. Group-level differences (*p <* 0.001 uncorrected, two-sided; cluster threshold: *p <* 0.05 with false discovery rate correction) were predominantly observed in the postcentral and precentral gyri. The complete list of clusters is presented in Table S1.

Additional analyses were performed correlating functional connectivity and ALS clinical metrics. These methods and results are presented in Supplementary Information. A meta-analysis of 57 ALS studies performed using NeuroQuery^84^, also presented in Supplementary Information, revealed activations in regions we found to be connected (i.e., somatosensory/motor cortex and cerebellum).

## 4 Discussion

In this data-driven, cross-sectional, ultra-high field rs-fMRI pilot study, we report significantly decreased functional connectivity between the superior sensorimotor cortex and bilateral cerebellar lobule VI in participants with ALS. *Post hoc* seed-to-voxel analyses of cerebellar lobule VI localize group-level differences in functional connectivity to bilateral precentral and postcentral gyri. While previous rs-fMRI studies have implicated the motor cortex and/or cerebellum in ALS, we are not aware of a previous report highlighting disruption in the specific link between the superior sensorimotor cortex and cerebellar lobule VI.

The upper motor neurons residing in the precentral gyri are well known to be involved in various disease mechanisms and neurodegeneration in ALS^17^. There is also growing evidence of the involvement of cerebellar structures in imaging and neuropathological studies in sporadic ALS, ALS/neurodegenerative syndromes arising due to C9orf72 or ataxin-2 repeat expansions and TBK1 mutations, and in primary lateral sclerosis (PLS)^85–88^. For example, pathological features associated with the production of C9orf72 transcripts with expanded repeats are the formation of nuclear ribonucleic acid foci in the frontal cortex, hippocampus, and cerebellum ^85^. Functional and diffusion neuroimaging studies support topographical mappings of cerebellar lobule VI to premotor and primary motor cortices reciprocally ^89^. Task-based fMRI has revealed that ocular and orofacial movements engage the medial aspect of cerebellar lobule VI^90–94^. During more complex motor tasks involving motor planning/learning or cognitive processing, activation patterns shift more laterally in lobules VI and VII^95,96^. Previous rs-fMRI studies have reported on such cortical-cerebellar circuits *in healthy volunteers* ^97–99^, further supporting the feasibility that this network is susceptible to disease-related impairment. The multifaceted role of cerebellar lobule VI therefore underscores the significance of its involvement in degeneration in ALS.

Previous studies have reported that cerebellar lobule VI and other regions of the cerebellum play a key role as a hub for diverse and integral motor and non-motor functions in ALS. A recent 3T study using an event-related motor paradigm reported the involvement of lobule VI in ALS participants with UMN predominant dysfunction^100^. A 3T diffusion tensor imaging (DTI) analysis of white matter tracts in ALS and PLS participants with pseudobulbar affect found abnormalities in the frontotemporal cortex, transverse pontine, and middle cerebellar peduncular circuits ^101^. A recent longitudinal 3T voxel-based morphometry study in sporadic ALS (excluding C9orf72 ALS participants) found significant cerebellar gray matter density reductions in multiple regions including bilateral cerebellar lobule VI that evolved on subsequent follow-up scans compared to matched controls ^102^. Another 3T DTI study showed widespread disruption of cerebellar white matter tracts along the dentato-rubro-thalamo-cortical and spino-cerebellar pathways in PLS that may be greater than ALS, implying the impact of longer duration motor neuron disease^103^. Such studies highlight the fact that studying the cerebellum and its neural connections is an important endeavor – yet remains challenging due to the relatively small size of many key anatomical structures.

The functional involvement of the cerebellum in ALS ^104^ has been reported in task-based ^41^,^42^,^105^,^100^ and resting state ^28,106,33,38^ fMRI studies at lower fields, and functional changes have been correlated to disease severity^106^, clinical variables ^100^, and task difficulty ^41^. However, participants with ALS have a reduced capacity for performing motor tasks associated with task-based fMRI, so rs-fMRI ^54,55,33,34,58,57,37,38,^ along with motor imagery paradigms^45^, are attractive options in studying diseases with motor implications. There is considerable variability in the literature on functional connectivity network patterns in ALS. Beyond motor task-related disability and heterogeneity in ALS participant characteristics, there are of course other challenges for developing and scaling fMRI as a biomarker for tracking disease progression in ALS including technological (e.g., scanner vendor, field strength) and methodological/statistical (acquisition, preprocessing choices, hypothesis-vs. data-driven approaches) considerations ^107,59.^

Earlier studies have reported abnormal connectivity/function involving the cerebellar and SMN areas in ALS ^28,41,38^. Specifically, 3T studies have highlighted the implications of ALS upon cerebello-cortical pathways. Both ALS participants and ALS mutation carriers have exhibited *increased* functional connectivity between the cerebellum and the precuneuscingulate-middle frontal resting state network compared to controls, which may indicate pre-symptomatic ALS cerebral pathology^33^. ALS participants have also shown decreased functional connectivity between the dentate nucleus in the cerebellum and both the supplementary motor area and left cerebellum lobule IV, which correlated positively with ALSFRS-R^38^. These alterations in functional connectivity, especially regarding cerebellar lobule IV and the primary motor cortex, may indicate alterations in the spino-cerebellar tract^38^. Furthermore, SMN correlations may increase in the earlier stage of disease^28^ and then decrease with more severe and longer duration disease^56,36^. An early 1.5T study reported decreased functional connectivity between the right SMN and right cerebellar lobule VI in ALS participants vs. healthy controls, but only in a subset of 16 ALS participants *with CST damage* (as measured via decreased fractional anisotrophy relative to controls and also ALS participants with undetectable CST damage) and only within the right hemisphere ^28^. A longitudinal rs-fMRI study (with 13 ALS and 3 PLS participants) observed functional connectivity decreases between the SMN and frontal pole and increases between the left fronto-parietal network and left primary motor cortex^36^. Another longitudinal 3T study (27 ALS participants vs. 56 controls) reported expansions in intrinsic motor, brainstem, ventral attention, and default mode/hippocampal networks that appeared to follow the same neuropathological pattern seen in Braak’s TDP43 staging in ALS ^34^. However, a cross-sectional study at 4.7T (20 ALS participants vs. 34 controls) used 10-mm diameter spheres within the pre-and post-central gyri and supplemental motor area to form a combined SMN time course and reported no significant group-level differences in the default mode network or SMN, or between subgroups of high and low UMN burden participants^56^.

Due to higher signal-to-noise and functional contrast-to-noise ratios, unique spatial contrasts, and increasing availability of 7T systems, high-resolution structural and functional imaging at 7T is poised to provide new insights into disease manifestation and progression ^108^ in ALS as well as other diseases. The primary advantages of 7T fMRI over lower fields is the greater-than-linear increase in BOLD sensitivity^109,110^ that permits smaller voxels without losing statistical power, and weighting toward smaller vessels supplying the superficial cortical structures, such as sensorimotor cortices, and smaller infratentorial tissue structures in close proximity to air-fluid-tissue interfaces such as cerebellum. One must keep in mind, however, that the mechanisms that give rise to an increase in BOLD contrast at higher fields also increase physiological noise ^111^, and thus care must be taken to mitigate confounding noise sources due to respiration, cardiac pulsatility, and subject motion. In this 7T study, the influence of physiological noise was modeled in CONN using ART-derived outliers, five principal components of anatomically-derived white matter and CSF segments, and the six estimated rigid-body motion parameters and their first derivatives. The appropriate impact of these regressors on each subject’s data was then reviewed using CONN’s suite of tools to visualize first-level fMRI data before and after denoising.

Our investigation has limitations that must, of course, be mentioned. First, this pilot study is a cross-sectional design with relatively small group sizes, so we are therefore not characterizing the trajectory of disease progression for individual participants but rather are performing a group-level comparison that includes inherent heterogeneity within the cohort of ALS participants. Second, while there is one identified C9orf72 positive ALS participant in this cohort, the genetic status and hence contribution of other non-genotyped ALS participants to the abnormal cerebellar connectivity results is unknown. Third, the fMRI data were acquired with a longer TR than what is commonly used nowadays, resulting in fewer degrees of freedom and limitations on how the analyses may be conducted. It is important to note that the high-resolution acquisition strategy was implemented with the methods available when this study commenced in mid-2011 to obtain fMRI data in a regime expected to have minimal physiological noise ^112,113^. Single-subject analyses were initially pursued in the analyses of these data, but the resultant degrees of freedom per subject after denoising necessitated the second-level analyses presented herein. A longer TR additionally precludes the study of higher frequency resting state signals – up to 0.8 Hz – which has become an important area of investigation in recent years ^114 –117^. Thus, future studies should use a TR of 1–1.5 sec and also acquire a longer resting state run to increase the fidelity of functional connectivity estimates ^118^. Finally, due to the high-resolution whole-brain acquisition strategy, the imaging FOV did not include the more caudal regions of the cerebellum; therefore, the results presented in this study cannot preclude the possible involvement of these regions of the cerebellum.

Reports of modulation in the cerebellum support the evidence that there are not only adaptive intracortical changes ^119,40,41,45,55,48,106,51,37,100^ in response to neurodegeneration in ALS, but also functional reorganization that occurs subcortically ^42,43,106,51,100^. In sum, these reports highlight the brain’s widespread plasticity and capacity to partially overcome cortical and spinal neuronal degeneration. (Our *post hoc* analysis of functional modulation restricted to the subcortex ^120^ may be found in Supplementary Information.) Future studies in ALS may employ recent acquisition advances, most notably simultaneous multi-slice imaging ^121^, to acquire high-resolution 7T fMRI data from the cerebrum and entire cerebellum with a reduced volume acquisition time. Accurate measurements of cerebellar volume should also be obtained to investigate possible relationships between changes in BOLD signals and gray matter atrophy. In addition to ALSFRS-R, the collection of neurocognitive clinical measures should be considered ^122^. Ultimately, larger cohorts and longitudinal studies at ultra-high field are necessary in various genetic subgroups and phenotypes of UMN-predominant motor neuron diseases to better understand the complex functional connectivity architecture from standpoints of both clinical relevance and tracking disease progression. Results from such studies may have a broader impact to inform the design of fMRI studies to better understand the involvement of various neural systems in other neurodegenerative diseases – including Alzheimer’s disease, Parkinson’s disease, Huntington’s disease, multiple sclerosis, frontotemporal dementia, and PLS – where the cerebellum is thought to be affected under the shadows of other key motor and cognitive neural pathways ^123,124,103.^

## 5 Conclusions

This pilot study acquired 7 Tesla resting state fMRI data in healthy controls and participants with ALS. An exploratory whole-brain analysis revealed a disruption in functional connectivity between the superior sensorimotor cortex (in the precentral gyrus) and bilateral cerebellar lobule VI. *Post hoc* analyses using atlas-derived left and right cerebellar lobule VI revealed decreased functional connectivity in ALS participants that predominantly mapped to bilateral postcentral and precentral gyri. Cerebellar lobule VI is associated with a wide range of key functions including complex motor and cognitive processing tasks. These findings add to the growing number of ALS reports implicating the cerebellum, and future studies are required with larger cohorts and known genetic status for all ALS participants.

## Acknowledgments

The authors acknowledge the generosity of our participants and their families. We also thank Dr. Alfonso Nieto-Castañón for helpful discussions on data preprocessing using the CONN toolbox. Imaging was performed at the Athinoula A. Martinos Center for Biomedical Imaging at the Massachusetts General Hospital using resources provided by the Center for Functional Neuroimaging Technologies (P41EB015896) and the Center for Mesoscale Mapping (P41EB030006), Biotechnology Resource Grants supported by the National Institute of Biomedical Imaging and Bioengineering, National Institutes of Health (NIH). The NIH also provided support through grants R00EB016689 and R01EB027779 (R.L.B.) and K23NS083715 (N.A). This research was also supported in part by the Harvard NeuroDiscovery Center, the Muscular Dystrophy Association, and the American Academy of Neurology (N.A.), and by the MGH/HST Athinoula A. Martinos Center for Biomedical Imaging. The content is solely the responsibility of the authors and does not necessarily represent the official views of the NIH.

