## Supplementary Information for "Ultra-high field (7T) functional magnetic resonance imaging in amyotrophic lateral sclerosis: a pilot study"

[1]Athinoula A. Martinos Center for Biomedical Imaging, Department of Radiology, Massachusetts General Hospital, Charlestown, MA, USA
[2]Department of Radiology, Harvard Medical School, Boston, MA, USA
[3]Harvard–Massachusetts Institute of Technology Health Sciences & Technology, Cambridge, MA, USA
[4]Sean M. Healey & AMG Center for ALS at Massachusetts General Hospital, Department of Neurology, Neurological Clinical Research Institute, Boston, MA, USA
[5]Department of Neurology, Harvard Medical School, Boston, MA, USA
[6]Department of Psychology, Northeastern University, Boston, MA, USA
[7]Department of Psychiatry, Massachusetts General Hospital, Boston, MA, USA
[8]Department of Brain and Cognitive Sciences, Massachusetts Institute of Technology, Cambridge, MA, USA
[9]Siemens Healthineers, Erlangen, Germany
[10]Mittelhessen University of Applied Sciences, Department of Life Science Engineering, Institute of Medical Physics and Radiation Protection, Giessen, Germany
[11]Spaulding Rehabilitation Hospital, Charlestown, MA, USA
[12]Department of Physical Medicine and Rehabilitation, Harvard Medical School, Boston, MA, USA
[13]Division of Neuroradiology, Massachusetts General Hospital, Boston, MA, USA
[14]Sanofi Genzyme, Cambridge, MA, USA
[†]These authors contributed equally to this work.
[‡]These authors share senior authorship.

**Table S1:** Summary of *post hoc* seed-to-voxel results using atlas-derived left, right, and bilateral cerebellar lobule VI as seed regions.

**Left lobule VI**

| MNI coordinates of cluster (and total # of voxels) | Brain regions within cluster | # of voxels and % coverage of region |
|---|---|---|
| –14 –34 +68 (1420) | Postcentral gyrus, right | 476 (15%) |
| | Postcentral gyrus, left | 337 (9%) |
| | Precentral gyrus, right | 179 (4%) |
| | Precentral gyrus, left | 109 (3%) |
| | Precuneus cortex | 36 (1%) |
| | Superior parietal lobule, right | 33 (2%) |
| | Superior parietal lobule, left | 32 (2%) |
| +30 –16 +54 (153) | Precentral gyrus, right | 116 (3%) |
| | Postcentral gyrus, right | 2 (< 1%) |

**Right lobule VI**

| MNI coordinates of cluster (and total # of voxels) | Brain regions within cluster | # of voxels and % coverage of region |
|---|---|---|
| +18 –46 +76 (523) | Postcentral gyrus, right | 238 (7%) |
| | Precentral gyrus, right | 209 (5%) |
| | Superior parietal lobule, right | 27 (2%) |

**Bilateral lobule VI**

| MNI coordinates of cluster (and total # of voxels) | Brain regions within cluster | # of voxels and % coverage of region |
|---|---|---|
| +20 –48 +78 (2068) | Postcentral gyrus, right | 695 (21%) |
| | Precentral gyrus, right | 550 (13%) |
| | Postcentral gyrus, left | 248 (7%) |
| | Superior parietal lobule, right | 133 (9%) |
| | Precentral gyrus, left | 97 (2%) |
| | Precuneus cortex | 28 (< 1%) |
| | Superior parietal lobule, left | 13 (1%) |
| | Lateral occipital cortex, right | 2 (< 1%) |

### Correlations between functional connectivity and clinical measures in ALS participants

Additional *post hoc* correlations with false discovery rate (FDR) correction were made between significant functional connectivity findings and clinical measures (and their subscales) in ALS participants. We computed the mean time series from all gray matter voxels within the cerebellum cluster in lobule VI to obtain one cerebellar time series per ALS participant. Likewise, we obtained one time series per subject from the bilateral motor cortex cluster. A connectivity value was calculated for each subject via Pearson's correlation coefficient of these two time series. Linear relations between these coefficients and clinical variables were then assessed.

Figure S1 presents correlations between clinical variables and functional connectivity in ALS participants. Although no significant associations were observed at the canonical level of $p < 0.05$ after FDR correction, connectivity was associated at a trend level ($p \sim 0.1$) with bulbar dysfunction (ALSFRS-R bulbar sub-domain) ($R = 0.46$, $R^2 = 0.21$, $p = 0.13$) and vital capacity ($R = 0.47$, $R^2 = 0.22$, $p = 0.12$). Power analyses suggest that these associations may have been significant ($p < 0.05$) with 18–19 participants, so these supplementary observations are therefore presented to support future studies investigating potential associations with larger participant cohorts.

### Connectivity impairments in ALS within the subcortex

Although assessing whole-brain connectivity impairments in ALS at 7T was the central aim of this study, we also performed a supplementary analysis probing functional connectivity impairments in ALS participants only within the subcortex.

Specifically, we used the Harvard-Oxford subcortical atlas[120] to parcellate preprocessed fMRI data into 16 subcortical regions (7 bilateral subcortical regions (thalamus caudate, putamen, pallidum, hippocampus, amygdala, and accumbens), mid-brain, and brainstem). Mean fMRI time series were obtained from each region in every subject. Functional connectivity was computed between all pairs of regions using Pearson's product-moment correlation coefficient, to provide a 16×16 connectivity matrix per subject. Group differences in functional connectivity are reported ($p < 0.05$, corrected for multiple comparisons using Bonferroni's method). This comparison was controlled for head motion (mean framewise displacement[83]).

We observed that functional connectivity between left caudate and brainstem was significantly lower in ALS participants compared to controls ($p = 0.00037$, $T = 3.69$, Cohen's $d = 1.64$). The MNI centroids of these regions were: caudate (–13, 10, 10), brainstem (0, –31, 40). Notably, the connectivity was largely negative in ALS participants ($\mu$ = –0.14, $\sigma$ = 0.13, range: –0.36 to +0.07) while in controls it was generally positive and closer to zero ($\mu$ = 0.07, $\sigma$ = 0.12, range: –0.12 to +0.20). This suggests that the connectivity between these two regions, which was generally positive and nearly uncorrelated in healthy controls, was generally negative and anticorrelated in ALS participants – implying that a dormant connection in health turned into a negative connection in ALS, with deactivation of brainstem related to activation of caudate and vice versa. Functional connectivity between right caudate and brainstem was also lower in ALS participants without multiple comparisons correction ($p = 0.0027$, $T = 2.38$, Cohen's $d = 1.08$).

Finally, Fig. S2 shows a significant association between ALS symptom severity (ALSFRS-R) and left caudate–brainstem connectivity ($R = -0.60$, $R^2 = 0.36$, $p = 0.041$). As noted earlier, such large negative connectivity was not observed in controls at all, indicating that large negative functional connectivity between left caudate and brainstem might be a biomarker[11] of ALS onset. This finding might also have applications in tracking disease progression if replicable in larger samples.

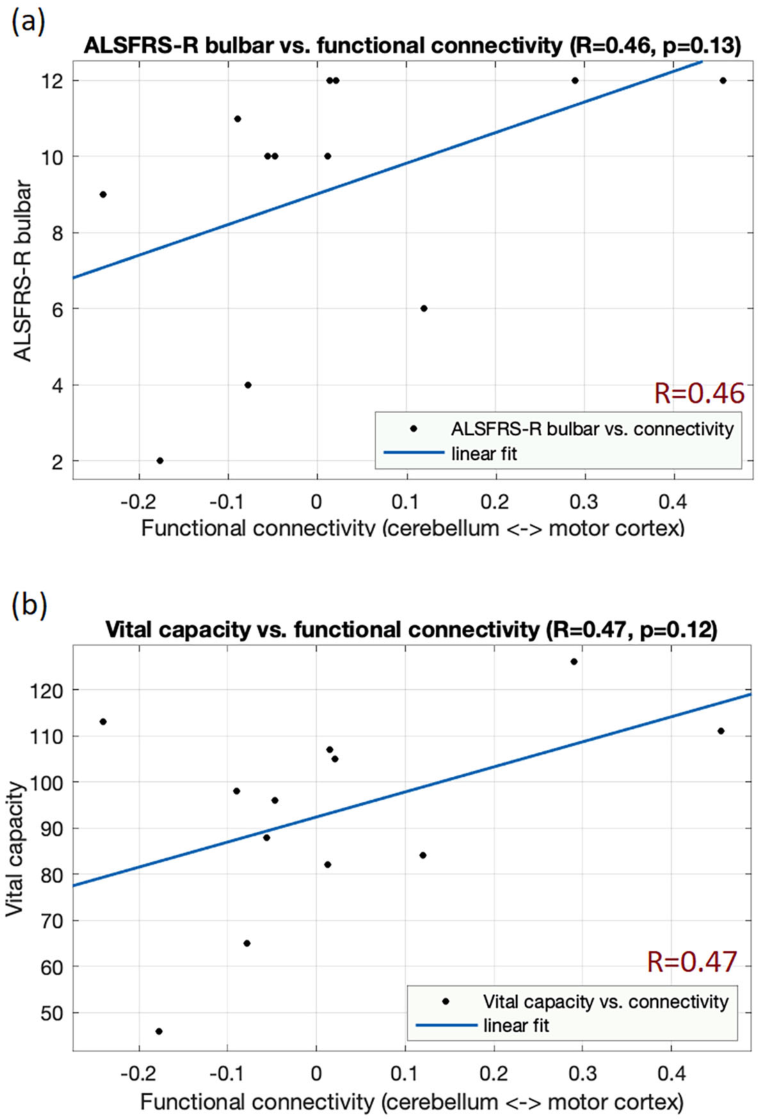


**Figure S1:** Associations between clinical variables and functional connectivity between cerebellum lobule VI and motor cortex. Although no significant associations were found after FDR correction, two associations exhibited trend-level significance: (a) ALSFRS-R bulbar subscale vs. connectivity ($R = 0.46$, $p = 0.13$) and (b) vital capacity vs. connectivity ($R = 0.47$, $p = 0.12$).

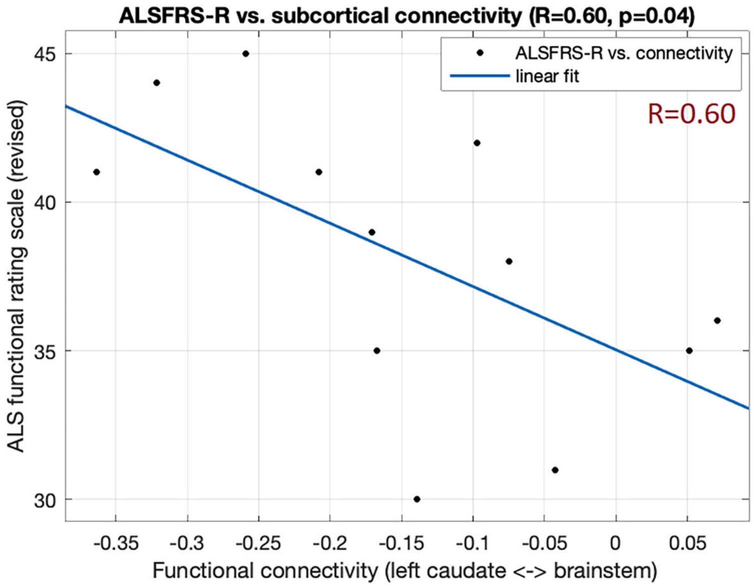


**Figure S2:** Association between ALS symptom severity (ALSFRS-R) and functional connectivity between left caudate and brainstem ($R = 0.60$, $p = 0.04$).

### Homoscedasticity when correlating functional connectivity and ALS clinical measures

To investigate the possibility of spurious associations with relatively small sample sizes, we also performed a quality check on the above three clinical associations. Linear regression makes the assumption of homoscedasticity – i.e., the variance of residuals is uniform across sample observations. Violation of this assumption, which can happen in low sample size regressions, renders the regression fit questionable. We tested the regression residuals of these associations for homoscedasticity using Engle's ARCH test, which tests the null hypothesis that the residuals exhibit no conditional heteroscedasticity ($p > 0.05$ implies that the residuals are homoscedastic and that the homoscedasticity assumption of linear regression is not violated). Two of the supplementary associations did not violate this assumption: (i) ALSFRS-R bulbar vs. connectivity ($p = 0.14$) and (ii) ALSFRS-R vs. subcortical connectivity ($p = 0.69$). One supplementary association did violate the assumption: vital capacity vs. connectivity ($p = 0.04$); as such, this association should be interpreted with caution.

### Correspondence between the results of this 7T study and a meta-analysis of ALS studies

NeuroQuery (neuroquery.org) is an fMRI-based meta-analysis tool that has gained popularity since its publication last year[84]. Meta-analysis activation likelihood estimate (ALE) maps show brain activations that are found to be most consistent across published fMRI activation studies. We used NeuroQuery to obtain ALE maps of ALS studies using the search term "amyotrophic lateral sclerosis" (https://neuroquery.org/query?text=amyotrophic+lateral+sclerosis). The results of this query (Fig. S3) show activations primarily in the somatosensory/motor cortex (±38, –22, 56), thalamus (±22, –22, 4), and cerebellum (±18, –54, –32). Our 7T results showed significantly impaired functional connectivity between two of the regions (i.e., somatosensory/motor and cerebellum) most consistently reported in previous ALS fMRI studies. We view the correspondence between the results presented herein and NeuroQuery's meta-analysis of 57 published ALS studies as further enhancing confidence in our 7T findings.

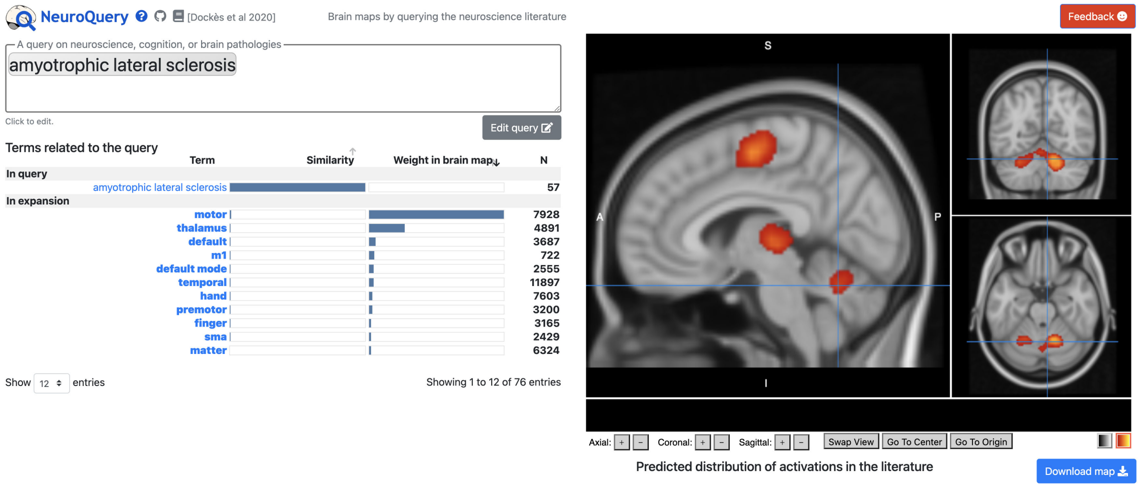


**Figure S3:** The output of a NeuroQuery[84] search, performed on February 12, 2021, using the search term "amyotrophic lateral sclerosis". The results of this meta-analysis of 57 published ALS studies show activation in the somatosensory/motor cortex, thalamus, and bilateral cerebellum.
